# RIP2 inhibits DOCK8-mediated platelet dense granule release and alleviates experimental myocardial infarction

**DOI:** 10.1101/2025.04.19.649674

**Authors:** Jianjun Zhang, Guanxing Pan, Yangyang Liu, Lin Chang, Liang Hu, Zhiyong Qi, Zhiling Zhang, Yan Zhang, Si Zhang, Jianlin Qiao, Xiaodong Xi, Kesheng Dai, Jianzeng Dong, Zhongren Ding

**Affiliations:** Department of Biochemistry and Molecular Biology, School of Basic Medical Sciences, Fudan University, Shanghai, China; Department of Cardiology, The First Affiliated Hospital of Zhengzhou University, Zhengzhou, China; Division of Cardiovascular Disease, Zhongshan Hospital, Fudan University, Shanghai, China; Department of Hematology, the affiliated hospital, Xuzhou Medical University, Xuzhou, China; Shanghai Institute of Hematology, Ruijin Hospital, Shanghai Jiaotong University School of Medicine, Shanghai, China; Jiangsu Institute of Hematology, Soochow University, Suzhou, China; 7Department of Clinical Laboratory, Shanghai Pudong Hospital, Fudan University Pudong Medical Center, Shanghai, China; School of Pharmacy, Department of Geriatrics of the Second Hospital, Tianjin Medical University, Tianjin, China

**Keywords:** RIP2, platelet, dense granule secretion, DOCK8, thrombosis, atherosclerosis, myocardial infarction

## Abstract

**Objective:** Receptor-interacting protein 2 (RIP2) is an essential mediator of inflammation and innate immunity downstream of pattern recognition receptors (PRRs). Platelets express RIP2, while its role in platelet activation, thrombosis, and myocardial infarction (MI) is unknown.

**Approach and Results:** Here we show that RIP2 deficiency enhances platelet dense granule secretion in response to GPIb and GPVI activation; platelet aggregation in whole blood and adhesion under arterial shear are also increased. Consistently, RIP2 inhibitor WEHI-345 potentiates human platelet dense granule secretion and inhibits RIP2 phosphorylation induced by thrombin and collagen. These phenotypes are translated into shorter bleeding time, accelerated FeCl_3_-induced arterial thrombosis. Importantly, platelets from patients with coronary artery disease and mice with atherosclerosis express lower RIP2, and RIP2 deficiency deteriorates MI and cardiac function in a mouse ischemia/reperfusion model. Mechanistically, we found that platelets express dedicator of cytogenesis protein 8 (DOCK8), which is sequestered by phosphorylated RIP2 (p-RIP2), causing inhibition of Cdc42 activation and subsequent dense granule release.

**Conclusions:** In conclusion, RIP2 inhibits platelet activation, thrombosis, and ameliorates MI. Our results also suggest that a novel PRR-independent pathway, p-RIP2-DOCK8-Cdc42, negatively regulates platelet activation downstream of GPIb and GPVI. Platelet RIP2 pathway may be promising for therapeutic intervention of atherothrombotic diseases from early atherosclerosis stage to arterial thrombosis and MI.

**Highlights:**

- RIP2 inhibits platelet dense granule release NOD2-independenly via pRIP2-DOCK8-Cdc42.
- The negative regulation of RIP2 on platelet activation is translated into enhanced thrombosis, exacerbated myocardial infarction (MI).
- RIP2 pathway booster may be an approach for therapeutic intervention of atherothrombotic diseases.

## Introduction

Platelets play essential roles in hemostasis and atherothrombosis. In the vasculature with intact endothelial lining, platelets are maintained in quiescent state and circulate in the blood. Upon vascular injury, platelets are exposed to subendothelial matrix proteins including collagen and von Willebrand factor (vWF), which stimulate GPVI and GPIb-IX-V and trigger platelet adhesion, aggregation, granule secretion, and thromboxane A2 (TXA2) synthesis^1^. Soluble agonists including secreted ADP, synthesized TXA2, and locally generated thrombin further amplify platelet activation and stabilize hemostatic plug^2^. GPIb-IX-V also binds thrombin and plays an important role in platelet activation induced by low concentration thrombin^1^. To exert the physiological function of hemostasis, platelets must be tightly regulated and maintained in quiescent state in normal blood flow to prevent excessive activation and unwanted thrombosis^2^. Uncontrolled and excessive platelet activation as in the situation of atherosclerotic plaque rupture and erosion in the coronary artery will cause occlusive thrombosis, resulting in heart attack^3^, the leading cause of death worldwide. In addition to the critical role in atherothrombosis, platelets are also involved in the pathogenesis of atherosclerosis, the common underlying mechanism for coronary artery disease, ischemic stroke, and peripheral arterial disease^2, 4–10^.

The receptor-interacting protein kinase family comprising 7 members is a group of Ser/Thr protein kinases with a highly homologous amino terminal kinase domain and distinctly different carboxyl termini^10^. Receptor-interacting protein kinase 2 (RIP2), also known as RIPK2 or RICK, is an essential mediator of inflammation and innate immunity^11, 12^. It is widely accepted that RIP2 is downstream of pattern recognition receptors (PRRs) and is activated through homophilic CARD-CARD (Caspase Recruitment Domain) interactions with nucleotide-binding oligomerization domain (NOD) containing receptors including NOD2^13^. Although RIP2 is autophosphorylated during NOD2 activation^14^, increasing evidences suggest that RIP2 functions as a scaffolding protein with its kinase activity dispensable to mediate NOD2 signaling^15,16^.

Recently, another member of receptor-interacting protein family, RIP3, was shown to promote platelet activation and thrombosis^17^. We previously reported that platelets express RIP2, and RIP2 is phosphorylated in human platelets upon NOD2 receptor activation^18^; we also found that NOD2 activation potentiates platelet activation^18^. However, the role of RIP2 in platelet activation, thrombosis, and myocardial infarction is unknown. We hypothesized that RIP2, as a downstream effector of NOD2, may positively regulate platelet activation and thrombosis. Surprisingly, we found that RIP2 deficiency enhances platelet activation, thrombosis, and exacerbates myocardial infarction. Mechanistically, we revealed a noncanonical RIP2 signal pathway independent of PRRs.

## Materials and methods

Details on reagents, chemicals, human blood samples, platelet function (aggregation, ATP release, spreading, and clot retraction) studies are described in SI Appendix, SI Materials and Methods^18–20^.

### Animal studies

RIP2-/- mice were purchased from Jackson Laboratory (Bar Harbor, ME). Six-week-old male ApoE-deficient mice and C57BL/6J mice were purchased from Cavens (Changzhou, China). All animal procedures were performed according to the criteria outlined in the “Guide for the Care and Use of Laboratory Animals” prepared by the National Academy of Sciences and published by the National Institutes of Health (NIH publication 86-23 revised 1985). All animals were sacrificed by cervical dislocation under isoflurane anesthesia (5 vol%).

### Platelet serotonin assay

Murine washed platelets were stimulated with agonists under stirring in aggregometer as for aggregation assay. Platelet suspension was centrifuged at 600 g for 5 minutes at room temperature. The supernatant serotonin was determined using Serotonin Enzyme-Linked Immunosorbent Assay Kit (ELISA) from Abcam (Cambridge, UK) according to the manufacturer’s protocol.

### Whole blood platelet aggregometry

Whole blood aggregation study was performed using the impedance method as described in SI Appendix, SI Materials and Methods ^21^.

### Platelet flow cytometry studies

Mouse platelet GPIIb (CD41), GPIIIa (CD61), GPVI, GPIb (CD42b) expression, and α granule release (P-selectin expression) were assayed by flow cytometry as described in Appendix, SI Materials and Methods.

### Transmission electron microscopy

Washed murine platelets were fixed in 4% glutaraldehyde at 4℃ overnight and then centrifuged at 700 × g for 5 minutes. After washing with PBS and post-fixing with 2% osmium tetroxide for 1 hour, the platelet pellets were dehydrated and embedded in Epon. Ultrathin sections were stained with 2% uranyl acetate and lead citrate and observed with a Phillips CM-120 transmission electron microscope (Phillips Inc., Eindhoven, Netherlands).

### Platelet adhesion on immobilized collagen under flow conditions and FeCl_3_-injured thrombus formation in mouse mesenteric arteriole

Platelet adhesion on flow chamber^22^ and intravital microscopy of FeCl_3_-injured thrombus formation in mouse mesenteric arteriole^18, 23, 24^ were measured as previously reported and described in Appendix, SI Materials and Methods.

### Bleeding time assay

Mouse bleeding time was measured as by tail snipping as described previously with slight modification^25^. More details are available in Appendix, SI Materials and Methods.

### Platelet depletion/reconstitution model

Platelet depletion/reconstitution model was performed as described previously^18, 24^. Briefly, platelets of WT mice were depleted via intraperitoneal injection of rabbit anti-mouse thrombocyte serum 20 μL for 4 hours prior to reconstitution with platelets from WT or RIP2^-/-^ mice, followed by *in vivo* assay of FeCl_3_-induced thrombus formation.

### Mass spectrometry and active Cdc42 pulldown assay

See Appendix, SI Materials and Methods for detail.

### RT-PCR and real time PCR analysis of platelet RIP2 from patients with coronary artery disease and mice with atherosclerosis

Total RNA was extracted from platelets with Trizol reagent (Invitrogen, Carlsbad, CA). After RNA isolation, 1 μg of total RNA was reverse transcribed to cDNA using an RT-PCR kit (TaKaRa, Japan). Real time PCR was performed using specific primers (Table I in the online-only Data Supplement). The primers were synthesized by Sangon Biotech (Shanghai, China).

### Evaluation of aortic atherosclerotic lesions

ApoE^-/-^ mice on C57BL/6J background were fed high fat diet and high cholesterol diet for 12 weeks to induce atherosclerosis. Atherosclerotic lesions were evaluated as described in SI Appendix, SI Materials and Methods.

### Mouse ischemia/reperfusion myocardial infarction model

The myocardial infarction study was performed by an investigator blind to the animal assignments as previously describe^9, 26, 27^. Wild type mice with platelet depleted and reconstituted with WT or RIP2^-/-^ platelets were randomly assigned to myocardial infarction or sham groups, respectively. Detailed protocols are described in SI Appendix, SI Materials and Methods.

### Statistical analysis

All data are expressed as mean ± SEM. Data normality was determined by Shapiro-Wilk test. Student’s *t* test was used for comparisons between two groups. Unless otherwise stated, one-way or two-way ANOVA followed by Tukey post hoc analysis was used when more than 2 groups and variables were compared. Prism 7.0 (GraphPad Inc, San Diego, CA) and IBM SPSS Statistics 19.0 (IBM Co., Armonk, New York) were used for statistical analysis. *P* < 0.05 was considered to be statistically significant.

## Results

### RIP2 knockout enhances platelet dense granule secretion of washed platelets

We expected that RIP2 will also similarly promote platelet activation as RIP3^17^. We first studied the effect of RIP2 deficiency on platelet aggregation and dense granule secretion *in vitro* using lumi-aggregometer. Using washed platelets, we found that RIP2 deficiency did not significantly influence platelet aggregation in response to multiple agonists including thrombin, AYPGKF, ADP, collagen, collagen related peptide (CRP), botrocetin, and U46619. In contrast to our expectation, the simultaneously recorded ATP release from dense granule was strongly enhanced in RIP2 deficient (RIP2^-/-^) platelets in response to thrombin, collagen, CRP, and botrocetin (Figure 1A, D - F); ATP release induced by AYPGKF and U46619 was not influenced by RIP2 deficiency (Figure 1B and 1G). Consistent with our findings, oligophrenin1 (OPHN1) deficiency increased dense granule release without influence on platelet aggregation^28^. We recorded platelet aggregation and ATP release simultaneously, the tracings clearly showed that the dense granule release measured by ATP secretion initiated after aggregation, which may explain why aggregation was not enhanced by increased dense granule release in RIP2^-/-^ platelets, even at low agonist concentrations.

**Figure 1.**
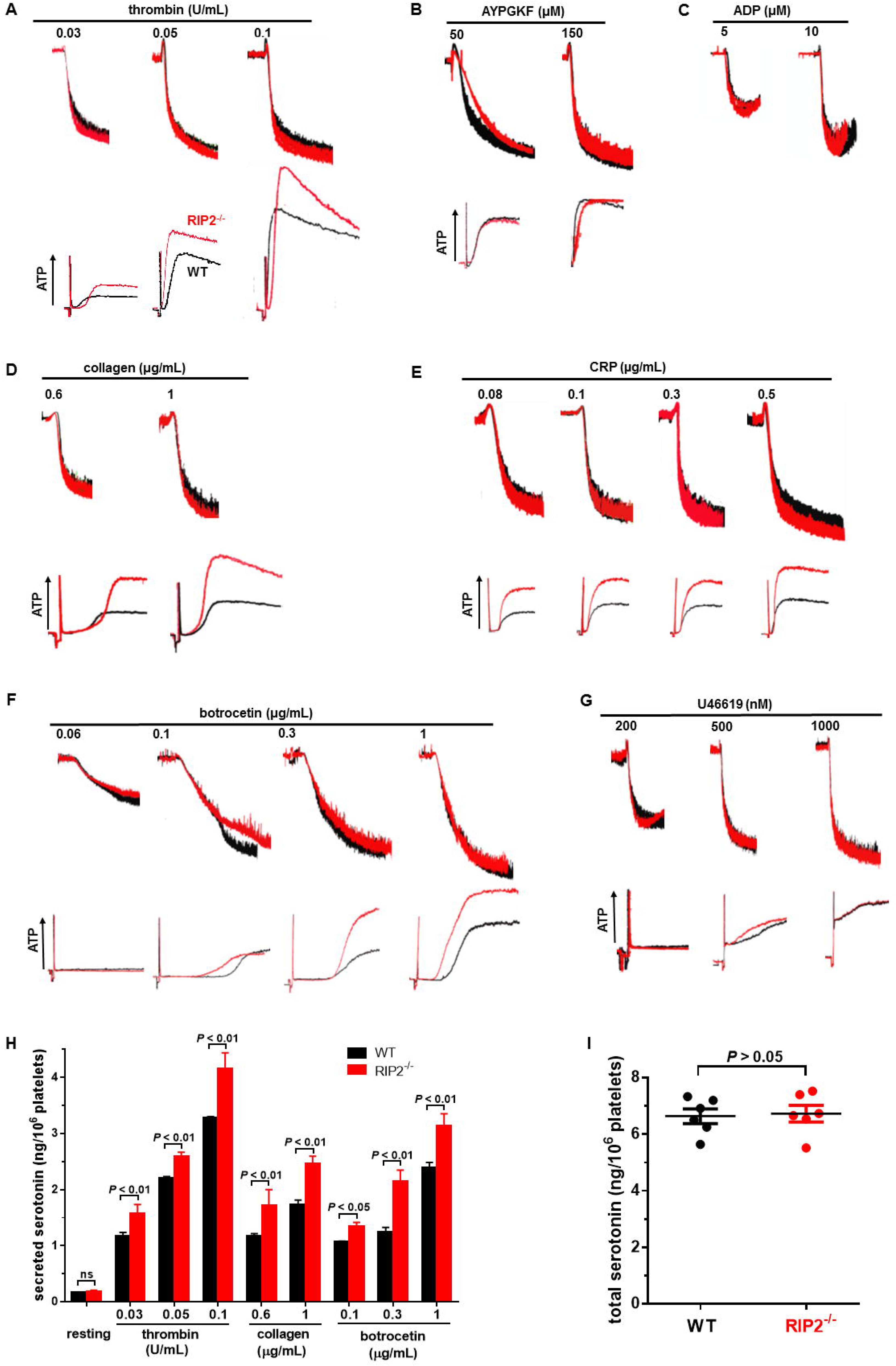
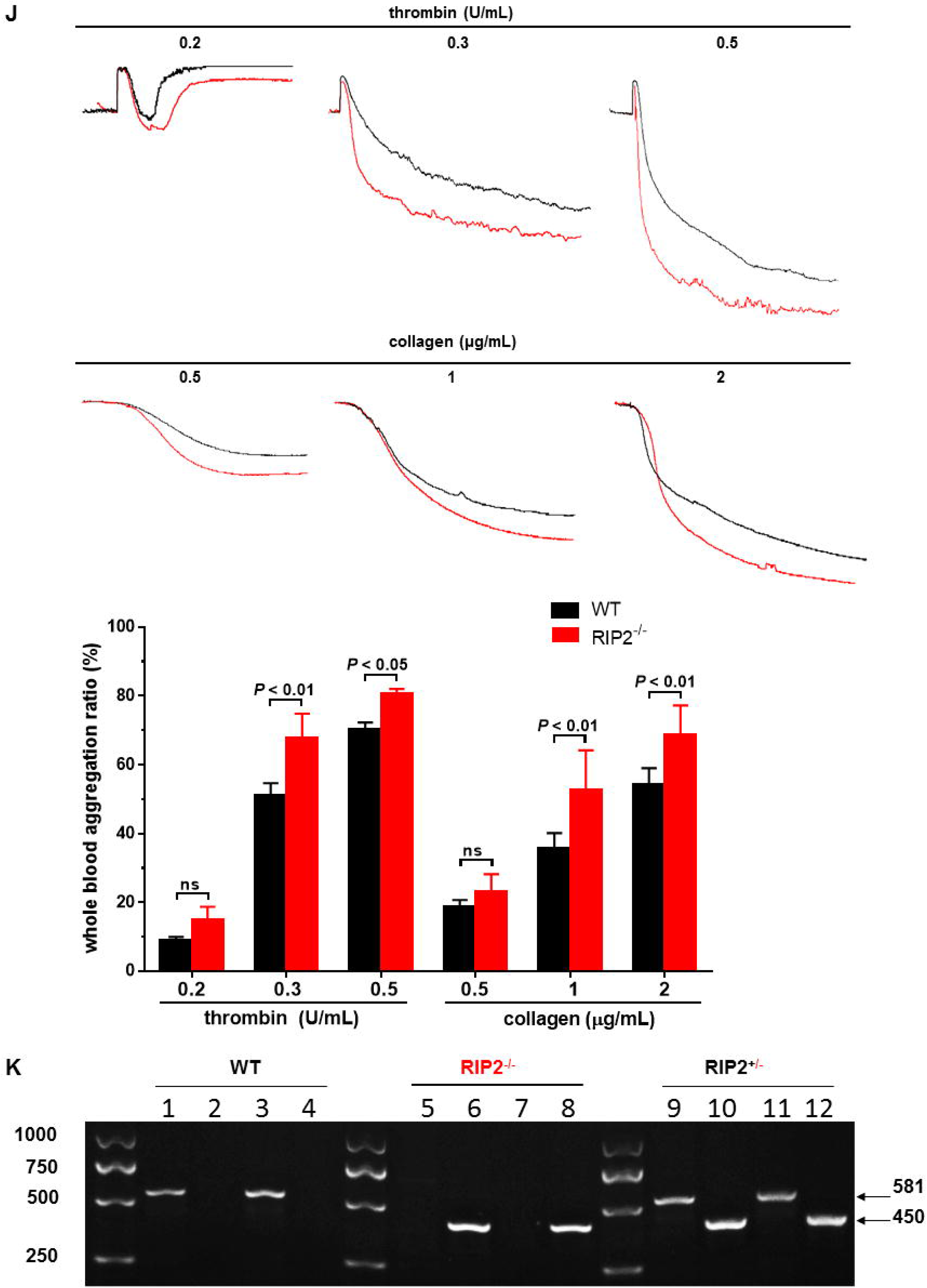
RIP2 knockout increases platelet dense granule release and whole blood platelet aggregation. Washed platelets from wild type (WT) and RIP2^-/-^ mice were stimulated with different concentrations of thrombin (**A**), AYPGKF (**B**), ADP (**C**), collagen (**D)**, CRP (**E**), botrocetin (**F**), and U46619 (**G**) at 37°C under constant stirring. Platelet aggregation and ATP release were simultaneously recorded using a Chrono-log lumi-aggregometer for 5 minutes. For botrocetin induced platelet activation, 6 μg/mL vWF was added. The tracings are representative of at least 5 independent experiments. **H**. RIP2 deficiency potentiates mouse platelet serotonin release in response to thrombin, collagen, and botrocetin. Serotonin in the supernatant of resting or activated platelets was assessed by ELISA. Data are expressed as means ± SEM (n = 6), two-way ANOVA followed by Bonferroni’s multiple comparisons test was used for statistical analysis. **I**. RIP2 deficiency does not change serotonin contents in mouse platelets. Murine washed platelets were lysed using equal volumes of lysis buffer on ice for 30 minutes. Serotonin was assessed in total platelet lysate. Data are expressed as means ± SEM (n = 6), Student’s t-test was used for statistical analysis. **J**. RIP2 knockout increases platelet in whole blood in response to thrombin and collagen. Platelet aggregation in whole blood was assayed using the impedance method. Whole blood 450 ml containing the same platelet counts from WT or RIP2^-/-^ mice were added to a cuvette containing a stir bar and 450 mL saline (warmed to 37°C prior to addition of blood). After equilibrating at 37°C for 10 minutes, thrombin or collagen was added to induce aggregation under stirring. Tracings were recorded using a Chrono-log lumi-aggregometer for 10 minutes. Representative tracings and the summary are (n = 6) are presented. Two-way ANOVA followed by Bonferroni’s multiple comparison test was used for statistical analysis. **K**. Genotyping of RIP2 knockout mice. The even-numbered lanes show 450 bp PCR products of mutant, whereas the odd-numbered lanes show the 582 bp PCR products of wild type. Primer information was provided in Table IV in the online-only Data Supplement.

We also measured serotonin release from platelet dense granule by ELISA. Consistent with dense granule ATP measured by lumi-aggregometry, RIP2 deficiency significantly potentiates platelet dense granule serotonin release in response to thrombin, collagen, and botrocetin without affecting total serotonin contents in mouse platelets (Figure 1H and I).

Platelet α granule release assayed by P-selectin expression in response to thrombin and AYPGKF is normal in RIP2^-/-^ platelets (Figure I in the online-only Data Supplement). In agreement with our findings, Yue et al reported that MINK1 deficiency only influenced platelet dense granule release without effect on platelet P-selectin release^29^.

### RIP2 deficiency augments platelet aggregation in whole blood

Whole blood platelet aggregometry replicates *in vivo* platelet activation conditions in the presence of white and red blood cells, and thus is more physiologically relevant to reflect platelet function^30^

We proceeded to measure platelet aggregation in whole blood of RIP2-deficent mice by the impedance method using Chrono-log aggregometer. As shown in Figure 1J, compared with wild type (WT) platelets, RIP2 deficiency significantly increased platelet aggregation in whole blood, which was not observed using washed platelets measured by light transmission aggregometry.

### RIP2 knockout does not change platelet dense granule, α granule contents and expression of major receptors

The enhanced dense granule secretion of RIP2^-/-^ platelets may be caused by increased dense granule contents. To address this issue, using transmission electron microscopy we checked platelet morphology. We found that RIP2^-/-^ platelets maintained normal platelet structure, dense and α granule numbers (Figure IIA and IIB in the online-only Data Supplement). Therefore, the enhanced dense granule secretion from RIP2^-/-^ platelets was not because of the change of dense granule number and contents. RIP2 deficiency slightly increased platelet count without influence on other hematologic parameters (Table II in the online-only Data Supplement) and expression of GPIIb (CD41), GPIIIa (CD61), GPVI, and GPIbα (CD42b) on platelets (Figure IIC and IID in the online-only Data Supplement).

### RIP2 knockout does not affect platelet spreading on fibrinogen and clot retraction

Platelet spreading is an early-phase outside-in signaling event downstream of platelet αIIbβ3 integrin activation^31^. Consistent with the normal aggregation of RIP2^-/-^ platelets, platelet spreading on fibrinogen was not affected (Figure IIIA in online-only Data Supplement). Similarly, clot retraction, the late-phase outside-in signaling event downstream of platelet αIIbβ3 integrin activation, also maintained normal in RIP2^-/-^ platelets compared with WT platelets (Figure IIIB in the online-only Data Supplement).

### RIP2 knockout increases platelet adhesion on collagen under arterial shear

To assess the contribution of platelet RIP2 to thrombus formation under physiological shear stress conditions, we investigated platelet adhesion to collagen using microfluidic whole blood perfusion assay at arterial shear rate. Whole blood from WT and RIP2^-/-^ mice was incubated with FITC labeled anti-mouse CD41 antibody to label platelets, perfused over collagen-coated BioFlux plates. As shown in Figure 2A, RIP2 deficiency significantly enhanced platelet adhesion on collagen at 1500 s^-1^ shear rate.

**Figure 2.** RIP2 deficiency increases platelet adhesion, hemostasis, and thrombosis. **A.** RIP2 deficiency increases platelet adhesion on collagen under arterial shear detected by microfluidics *in vitro*. Typical photomicrographs (original magnification, × 20) and summary showing increased platelet adhesion on collagen of whole blood from RIP2^-/-^ mice are provided, each dot presents the one experiment using blood from one mouse. Whole blood collected in 150 μM PPACK was fluorescently labeled by incubation with FITC labeled anti-mouse CD41 antibody for 30 minutes, and then perfused through collagen coated BioFlux plates at a shear rate of 1500 s^-1^ for 5 minutes. Student’s t test was used for statistical analysis (mean ± SEM, n = 6). **B.** Shorter bleeding time in RIP2^-/-^ mice than WT mice. Two-tailed Mann-Whitney test was used for statistical analysis (mean ± SEM, n = 18). **C**. Representative images showing increased thrombus formation in mesenteric arterioles of RIP2^-/-^ mice. The right is the summary showing dramatically less complete occlusion time of mesenteric arterioles of RIP2 mice (mean ± SEM, n = 8). **D.** Representative images showing increased thrombus formation in mesenteric arterioles of WT mice repopulated with platelets from RIP2^-/-^ donor mice. The right is summary showing significantly less complete occlusion time of mesenteric arterioles of WT mice repopulated with RIP2^-/-^ platelets (mean ± SEM, n = 8). Calcein was used to label platelets. Thrombosis was induced by FeCl_3_ injury and recorded with intravital microscopy. Time after FeCl_3_ injury is indicated at the top of each image. Two-tailed Mann-Whitney test was used for statistical analysis.

### Increased *in vivo* hemostasis and arterial thrombosis in RIP2^-/-^ mice

To explore whether the enhanced platelet activation *in vitro* can be translated into increased hemostasis and thrombosis *in vivo*, we first measured tail snip bleeding time to determine the role of RIP2 in hemostasis. As expected, we observed obviously shorter bleeding time in RIP2^-/-^ mice compared to the WT littermates (Figure 2B), indicating enhanced hemostasis in RIP2^-/-^ mice. Consistently, using intravital fluorescence microcopy we observed exaggerated thrombus formation after FeCl_3_ injury to mesenteric arterioles in RIP2^-/-^ mice (Figure 2C).

To evaluate whether the RIP2 in platelets specifically play the critical role in thrombus formation, using the platelet depletion/reconstitution model to create the chimeric mice lacking platelet RIP2 only, we found that the loss of platelet RIP2 did accelerate FeCl_3_-induced thrombus formation in mouse mesenteric arterioles (Figure 2D). In line with our findings, Levin et al found that mice transplanted with RIP2^-/-^ bone marrow unexpectedly displayed more severe atherosclerosis despite decreased inflammation^32^.

### RIP2 inhibitor WEHI-345 potentiates platelet dense granule release

RIP2 undergoes autophosphorylation at serine 176 upon activation of upstream PRRs^18, 33^, and p-RIP2 (Ser 176) can be used to reflect RIP2 activation state^33^. WEHI-345 is a potent and selective inhibitor of RIP2 with IC_50_ of 130 nM^34^. At 10 nM, we found that WEHI-345 potentiated dense granule release of human platelets induced by thrombin and collagen (Figure 3A), consistent with the heightened dense granule release from RIP2^-/-^ platelets (Figure 1A and 1D). When p-RIP2 (Ser 176) was checked, we found that 10 nM WEHI-345 significantly inhibited RIP2 phosphorylation induced by thrombin and collagen (Figure 3B). We further examined the effects of RIP2 inhibition on dense granule using mouse platelets. At 100 nM, WEHI-345 potentiated thrombin and collagen induced dense granule release of platelets from WT but not RIP2^-/-^ mice (Figure 3C and 3D), further confirming the negative regulation of RIP2 on platelet dense granule secretion; these results also suggest that the negative regulation of RIP2 on platelet dense granule release is mediated by RIP2 phosphorylation.

**Figure 3.**
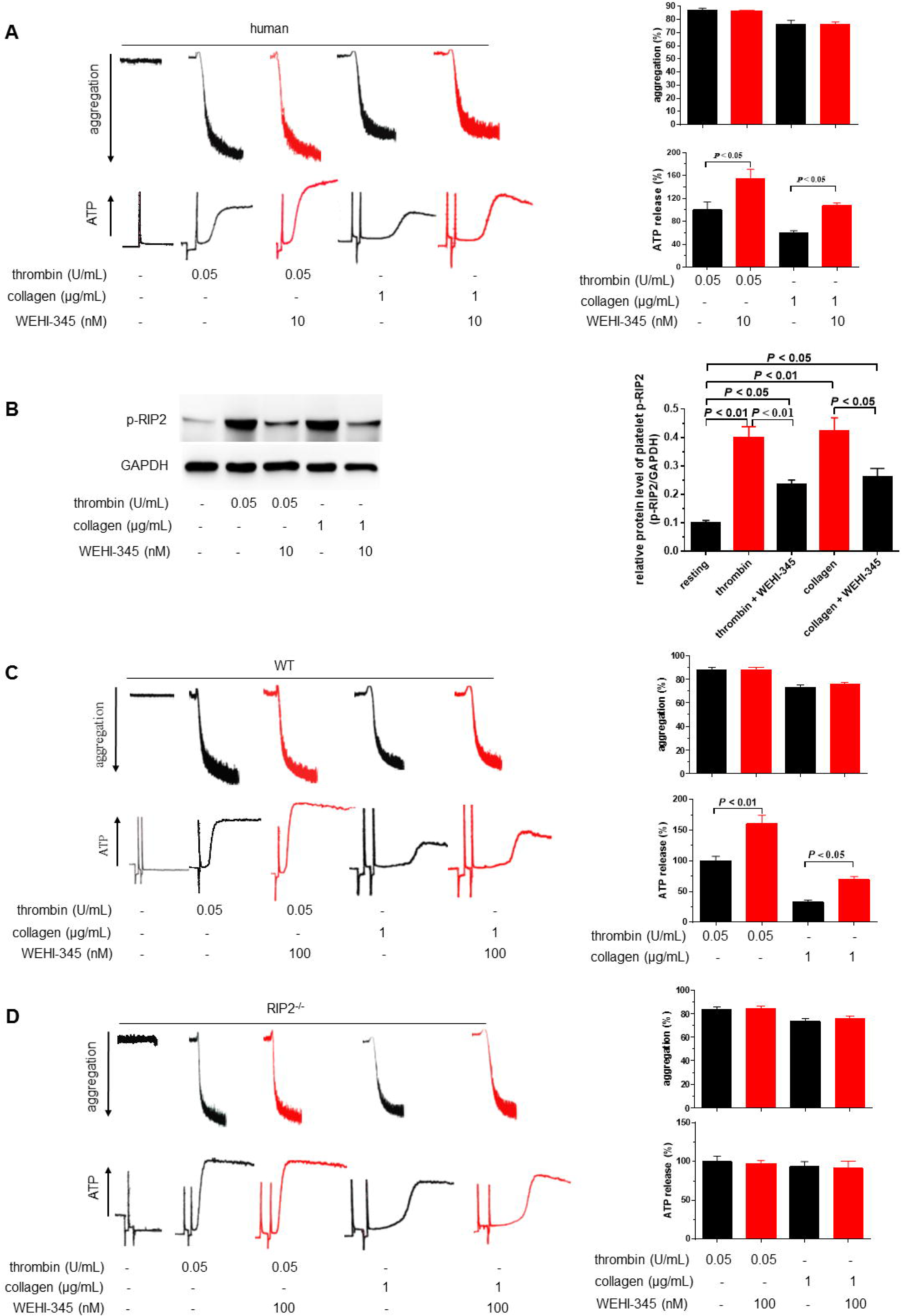
RIP2 inhibition potentiates dense granule release in human and mouse platelets stimulated with thrombin and collagen. **A**. RIP2 inhibitor WEHI-345 spotentiates ATP release without influence on aggregation of human platelets induced by thrombin and collagen. Human washed platelets were incubated with RIP2 inhibitor WEHI-345 or vehicle at 37°C for 20 min, thrombin or collagen was then added to initiate platelet aggregation and ATP release, recorded simultaneously with aggregometer under stirring. Representative tracings and summary of 5 independent experiments were presented (mean ± SEM. ATP release is normalized as ATP release at thrombin 0.05 U/mL taken as 100%). **B.** RIP2 inhibitor WEHI-345 suppresses RIP2 phosphorylation induced by thrombin and collagen. Platelet samples from panel A were analyzed by Western blot with anti-phospho-RIP2 (Ser 176) antibody. Results shown are representative of 5 independent experiments and the summary (mean ± SEM). Unpaired t test was used for statistical analysis. **C** and **D.** RIP2 inhibitor WEHI-345 potentiates ATP release of wild type (**C**), but not RIP2^-/-^ (**D**) mouse platelets induced by thrombin and collagen. Simultaneously recorded aggregation was not influenced by WEHI-345 pretreatment for 20 minutes. Representative tracings and summary of 5 independent experiments were presented. One-way ANOVA followed by Tukey post hoc analysis was used for statistical analysis.

Classically, RIP2 is regarded as an essential mediator relaying signal transduction downstream of PRR activation. RIP2 deficiency and inhibition potentiate platelet activation induced by thrombin, collagen, and CRP (Figure 1) suggests that RIP2 may also function downstream of GPIb and GPVI activation independently of PRRs. In line, thrombin and collagen induce RIP2 phosphorylation in the absence of PRR activation (Figure 3B and 4A). Moreover, NOD2 deficiency does not impair platelet RIP2 phosphorylation induced by thrombin and collagen (Figure IV online-only Data Supplement), further supporting NOD2-independent RIP2 activation.

**Figure 4.**
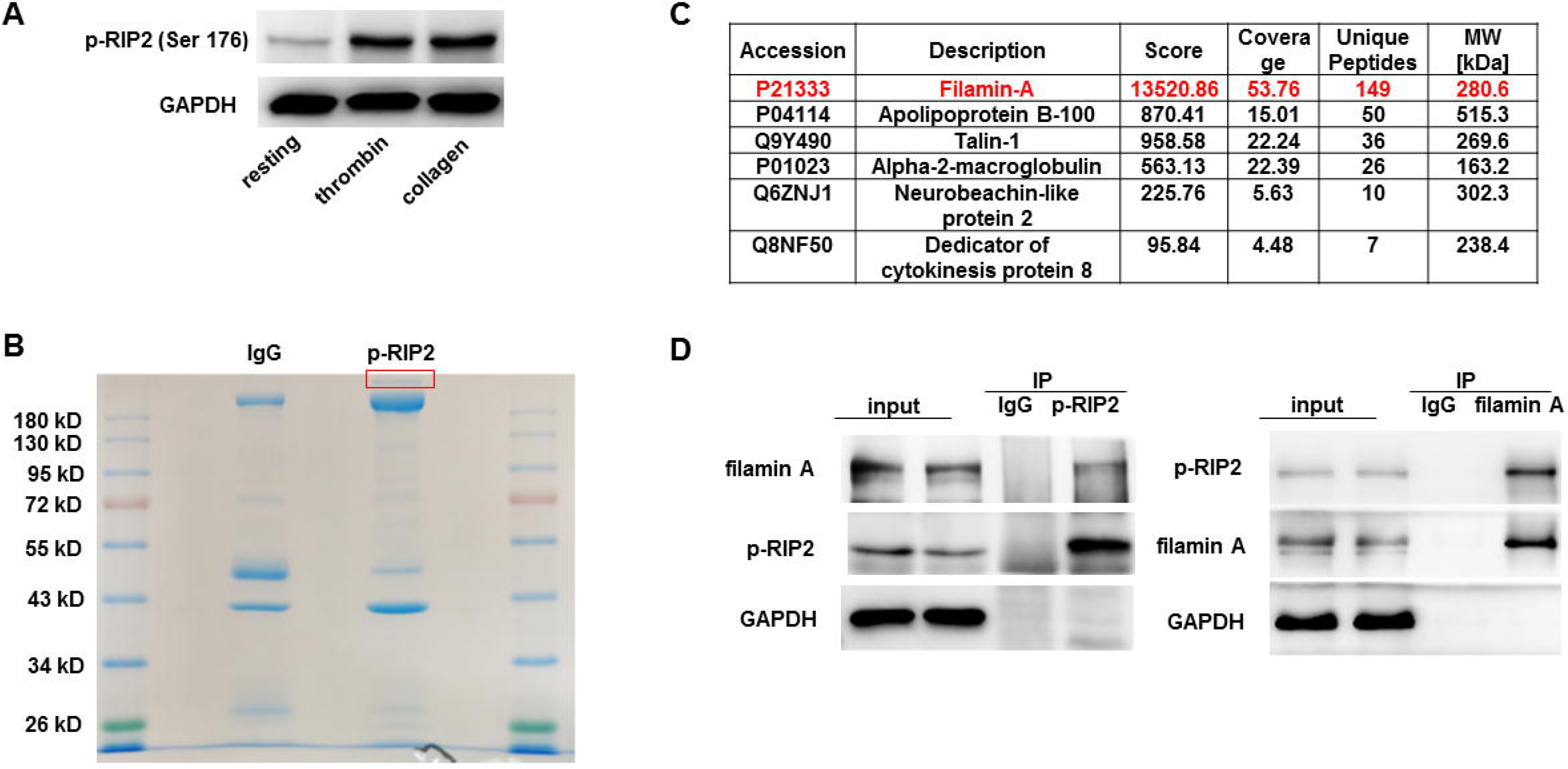
p-RIP2 interaction with filamin A identified by mass spectrometry and co-immunoprecipitation. **A**. Thrombin and collagen induce phosphorylation of RIP2 at serine 176, p-RIP2 (Ser 176), in human platelets. Human washed platelets were stimulated with thrombin 0.05 U/mL or collagen 1 µg/mL under stirring for 3 minutes, p-RIP2 (Ser 176) was measured by Western blot. Results shown are representative of 3 experiments using platelets from different donors. **B.** Coomassie blue staining of the immunoprecipitated proteins separated by 10% SDS-PAGE. Lysates from human platelets stimulated with thrombin were immunoprecipitated with antibody against p-RIP2 (Ser 176) or rabbit IgG as a control. A p-RIP2 specific band above 180 kDa specifically binding p-RIP2 (highlighted in red square) were sliced, in-gel trypsin digested, and subjected to mass spectrometry for protein identification. **C**. The p-RIP2 specific band in panel B was identified to be 280 kDa filamin A by mass spectrometry. **D.** Immunoblot after immunoprecipitation further confirmed the interaction between p-RIP2 and filamin A in thrombin stimulated human platelets. Immunoprecipitation of filamin A or p-RIP2 from human platelets extracts, using antibody against p-RIP2 (Ser 176) (left) or filamin A (right). Lane input is 10% input. Western blot analysis was performed using monoclonal antibody against filamin A or p-RIP2. Results shown are representative of at least 3 independent experiments.

### p-RIP2 binds both filamin A and GPIbα, RIP2 knockout promotes GPIbα dissociation from filamin A

Given the finding that RIP2 deficiency and p-RIP2 (Ser 176) inhibition potentiates platelet dense granule secretion induced by thrombin and collagen, we set out to dissect the underlying mechanism. We found that both thrombin and collagen induced phosphorylation of RIP2 (Ser 176) in human platelets (Figure 4A). In thrombin-stimulated human platelets, a 280 kDa protein coimmunoprecipitated with p-RIP2 (Ser 176) was identified as filamin A by mass spectrometry (Figure 4B and 4C). The interaction between p-RIP2 and filamin A was further confirmed by immunoblotting filamin A and p-RIP2 after immunoprecipitating p-RIP2 and filamin A, respectively (Figure 4D). We did not find the interaction between RIP2 and filamin A using the same methods (data now shown).

We further showed that p-RIP2 also bound to GPIbα (Figure 5A), and RIP2 knockout impaired GPIbα binding to filamin A in both resting platelets and platelets activated by thrombin and collagen (Figure 5B and 5C), suggesting that p-RIP2 strengthens the association between GPIbα and filamin A by binding both. Importantly, we noticed that thrombin and collagen stimulation promoted GPIbα dissociation from filamin A (Figure 5B and 5C). We thus proposed that p-RIP2 strengthens GPIbα binding filamin A by binding to both, constrains platelets in resting state, and prevents overactivation upon stimulation.

**Figure 5.**
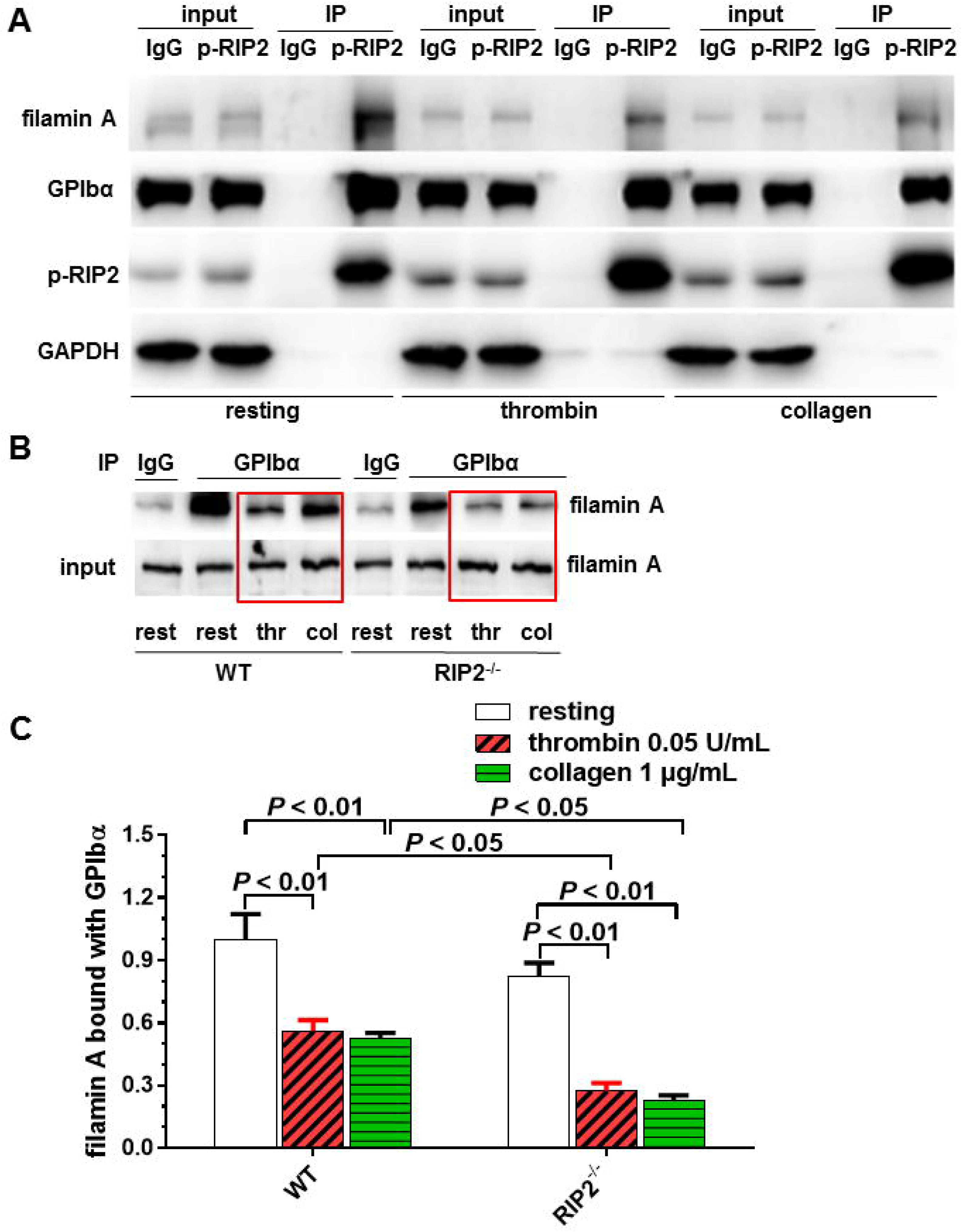
p-RIP2 binds both GPIbα and filamin A, RIP2 knockout promotes GPIbα dissociation from filamin A. **A.** p-RIP2 associates with both GPIbα and filamin A in resting and activated human platelets. Human platelets were activated by 0.05 U/mL thrombin or 1 µg/mL collagen, interaction between p-RIP2 and GPIbα or filamin A was detected by immunoprecipitation with antibody against p-RIP2 (Ser 176) from the lysates of platelets. A nonspecific rabbit IgG was used a control. Results shown are representative of at least 3 experiments using platelets from different donors. **B.** Decreased GPIbα-filamin A association in activated mouse platelets; RIP2 deficiency attenuates GPIbα-filamin A association in both resting and activated platelets. Mouse platelets were stimulated with 0.05 U/mL thrombin (thr) or 1 µg/mL collagen (col), GPIbα was immunoprecipitated from platelet lysates using a rat anti-mouse GPIbα antibody and filamin A was detected by Western blot. A nonspecific rat IgG was used a control. Results shown are representative of 5 experiments using platelets from different mice. **C.** Summary of panel B (mean ± SEM, n = 5). Two-way ANOVA followed by Tukey post hoc analysis was used for statistical analysis. Results were quantified as the ratio of the GPIbα bound filamin A normalized to the resting value of WT platelets taken as 1.

### Accelerated Cdc42 activation contributes to the enhanced dense granule release in RIP2^-/-^ platelets

Small GTPase Cdc42 has been reported to regulate platelet dense granule release, though opposite results have been reported^35, 36^. Actually, Cdc42 has been reported to bind filamin A^37^ and suggested to be a GPIb-specific downstream signaling molecule^35^. After showing p-RIP2 binds both GPIbα and filamin A (Figure 5A), we sought to explore the role of Cdc42 in the negative regulation of RIP2 on platelet dense granule release. We examined Cdc42 activity in RIP2^-/-^ platelets. Compared with WT mouse platelets, we observed significantly stronger Cdc42 activity as measured by GTP bound Cdc42 in RIP2^-/-^ platelets under both resting and activated conditions stimulated by thrombin and collagen (Figure 6A). We think that the enhanced Cdc42 activity contributes to the increased dense granule release in RIP2^-/-^ platelets in response to thrombin, collagen, CRP, and botrocetin (Figure 1).

**Figure 6.**
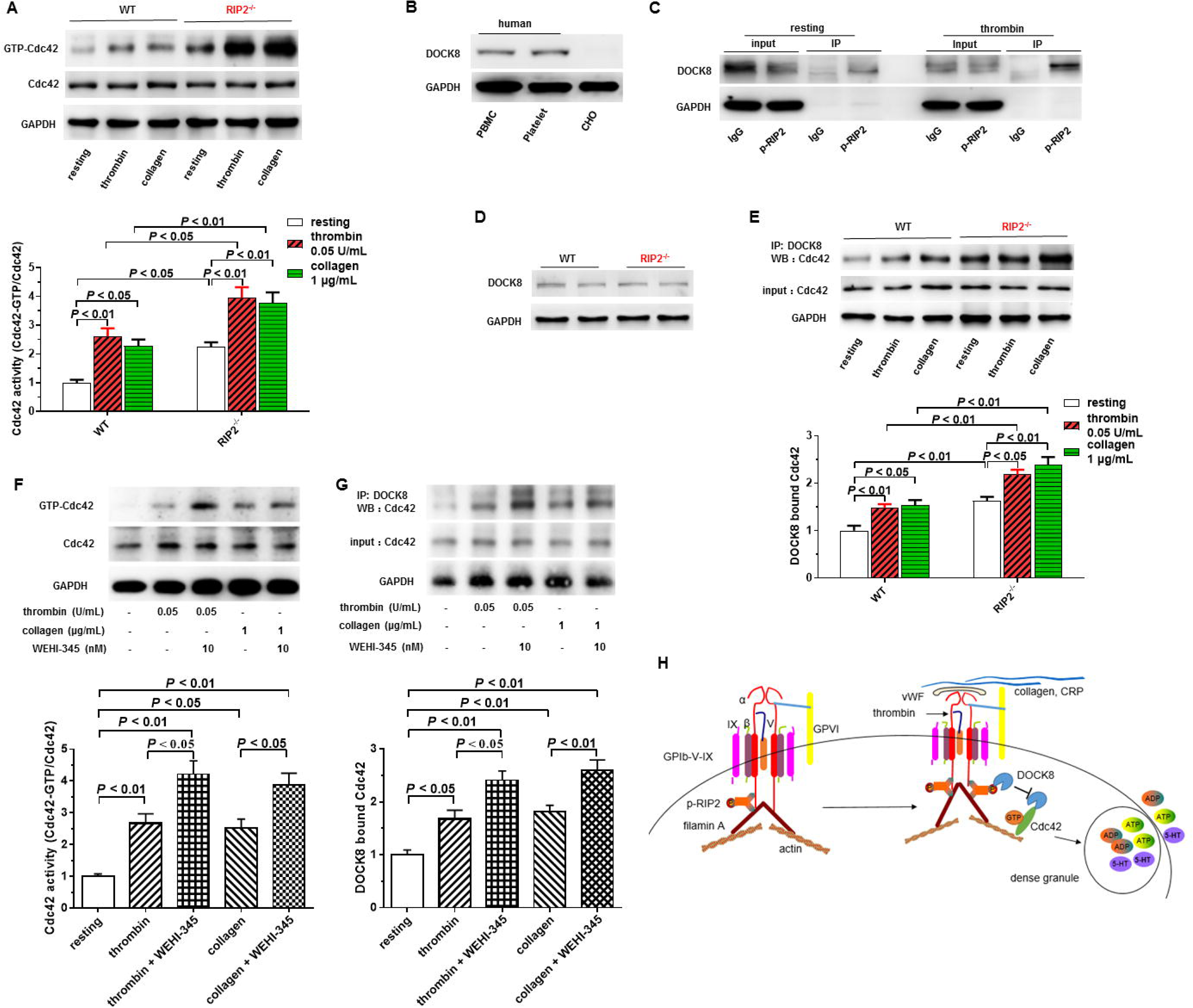
RIP2 deficiency increases DOCK8 mediated Cdc42 activation in platelets. **A**. Increased Cdc42 activation in platelets from RIP2^-/-^ mice as evidenced by enhanced GTP bound Cdc42. Platelets were stimulated with thrombin and collagen for 60 and 90 seconds, respectively, under stirring in an aggregometer at 37°C. Platelets lysates were subjected to pulldown assays using the GST-fusion Cdc42-binding domain of PAK1 before immunoblotting with anti-Cdc42 antibody. Upper panel is representative of 6 independent experiments using platelets from different mice with similar results. Bottom panel is the summary quantified as the ratio of the GTP-bound Cdc42 to the total Cdc42 normalized to the resting value of WT platelets taken as 1. Data are expressed as mean ± SEM (n = 6). Two-way ANOVA followed by Tukey post hoc analysis was used for statistical analysis. **B.** DOCK8 expression in human platelets detected by Western blot. PBMC (peripheral blood mononuclear cell) and CHO (Chinese hamster ovary cell) cells were used as positive control and negative control, respectively. **C.** DOCK8 associates with p-RIP2 in both resting and activated human platelets stimulated by thrombin. Human washed platelets were stimulated with vehicle or thrombin 0.05 U/mL for 3 minutes as in Figure 4A; DOCK8 association with p-RIP2 was detected by co-immunoprecipitation. **D.** Mouse platelets express DOCK8 at protein level, and RIP2 deficiency does not affect DOCK8 expression. Results shown in panel B - D are representative of at least 3 different experiments. **E.** Increased interaction between DOCK8 and Cdc42 in RIP2 knockout platelets. Platelets were stimulated as in panel A and B, DOCK8-bound Cdc42 was measured by immunoprecipitation of DOCK8 followed by Western blot detection of Cdc42. Results are representative of 7 experiments using platelets from different mice. Typical immunoblot and the summary are provided. Data were quantified and analyzed as in panel A (n = 7). **F.** RIP2 inhibitor WEHI-345 increases Cdc42 activity in human platelets as evidenced by enhanced GTP bound Cdc42. Typical immunoblot representative of 6 independent experiments using platelets from different donors and the summary are provided. Human washed platelets were incubated with RIP2 inhibitor 10 nM WEHI-345 or vehicle at 37°C for 20 min, and then stimulated with thrombin and collagen for 60 and 90 seconds, respectively, under stirring in an aggregometer at 37°C. Platelets lysates were subjected to pulldown assays using the GST-fusion Cdc42-binding domain of PAK1 before immunoblotting with anti-Cdc42 antibody. Data were quantified and analyzed as in panel A. One-way ANOVA followed by Tukey post hoc analysis was used for statistical analysis. **G.** Increased interaction between DOCK8 and Cdc42 in human platelets treated with RIP2 inhibitor WEHI-345. Platelets were treated as in panel F, DOCK8-bound Cdc42 was measured by immunoprecipitation of DOCK8 followed by Western blot detection of Cdc42. Representative immunoblot and the summary 6 experiments are presented. Data were quantified and analyzed as in panel F (n = 6). **H.** Proposed model depicting that p-RIP2 binds with both GPIbα and filamin A, strengthening the association between GPIbα and filamin A, inhibiting platelet activation downstream of GPIbα. Specifically, phosphorylated RIP2 binds and sequesters DOCK8, prevents DOCK8 interaction with Cdc42, inhibits Cdc42-induced dense granule secretion.

The role of Cdc42 in platelet dense granule is inconsistent^35, 36^. Our finding that RIP2 deficiency enhances dense granule release and Cdc42 activity is consistent with the presumed function of Cdc42, namely, it positively regulates platelet dense granule secretion^36^. Notably, Cdc42 deficient platelets were reported to fully spread on fibrinogen^35^; this is in line with our findings that RIP2 deficiency does not affect platelet spreading (Figure IIIA in online-only Data Supplement), though dense granule release is enhanced.

### Increased DOCK8-Cdc42 interaction mediates the elevated Cdc42 activation in RIP2^-/-^ platelets

DOCK8 (dedicator of cytogenesis protein 8) is a guanine nucleotide exchange factor for Cdc42 activation^38^. Our mass spectrometry revealed an interaction between p-RIP2 and DOCK8 in thrombin-stimulated human platelets (Figure 4C), suggesting DOCK8 expression in platelets; DOCK8 expression in human platelets and its interaction with p-RIP2 were further confirmed by Western blot and immunoprecipitation (Figure 6B and 6C). Using Western blot, we recapitulated the expression of DOCK8 in mouse platelets, and found that RIP2 deficiency does not affect DOCK8 expression (Figure 6D). We speculated that DOCK8 may be involved in the elevated Cdc42 activation in RIP2^-/-^ platelets. We next asked whether RIP2 deficiency affected DOCK8 interaction with Cdc42. As expected, we detected significantly higher DOCK8-bound Cdc42 in RIP2^-/-^ platelets under both resting and activated conditions stimulated by thrombin and collagen (Figure 6E), coinciding with the significantly increased Cdc42 activity in RIP2^-/-^ platelets (Figure 6A). In line, RIP2 inhibitor WEHI-345, significantly increased Cdc42 activity (Figure 6F) and DOCK8-bound Cdc42 (Figure 6G) in human platelets in response to thrombin and collagen. We thus proposed that RIP2 negatively regulates platelet dense granule secretion by inhibiting Cdc42 activation as a result of impaired DOCK8 activation on Cdc42 (Figure 6H).

### Impaired platelet RIP2 expression in patients with coronary artery disease and ApoE^-/-^ mice with atherosclerosis

Promoted by the negative regulation RIP2 in platelet activation and experimental arterial thrombosis, we investigated its clinical significance in atherothrombotic disease. Consistently, we found that platelets from patients with coronary artery disease expressed significantly less RIP2 at protein level compared with heathy subjects (Table III in the online-only Data Supplement), though both groups have similar platelet RIP2 mRNA level (Figure 7A and 7B). These findings were recapitulated in platelets from ApoE^-/-^ mice with atherosclerosis: platelets from ApoE^-/-^ mice with atherosclerosis expressed significantly less RIP2 protein while maintaining relatively normal mRNA level (Figure 7C - E), in line with the anti-atherosclerotic role of RIP2 reported by Levin et al^32^. The impaired platelet RIP2 expression may dampen its negative regulation on platelet activation and contribute to the pathogenesis of atherosclerosis and atherothrombosis of coronary artery disease.

**Figure 7.**
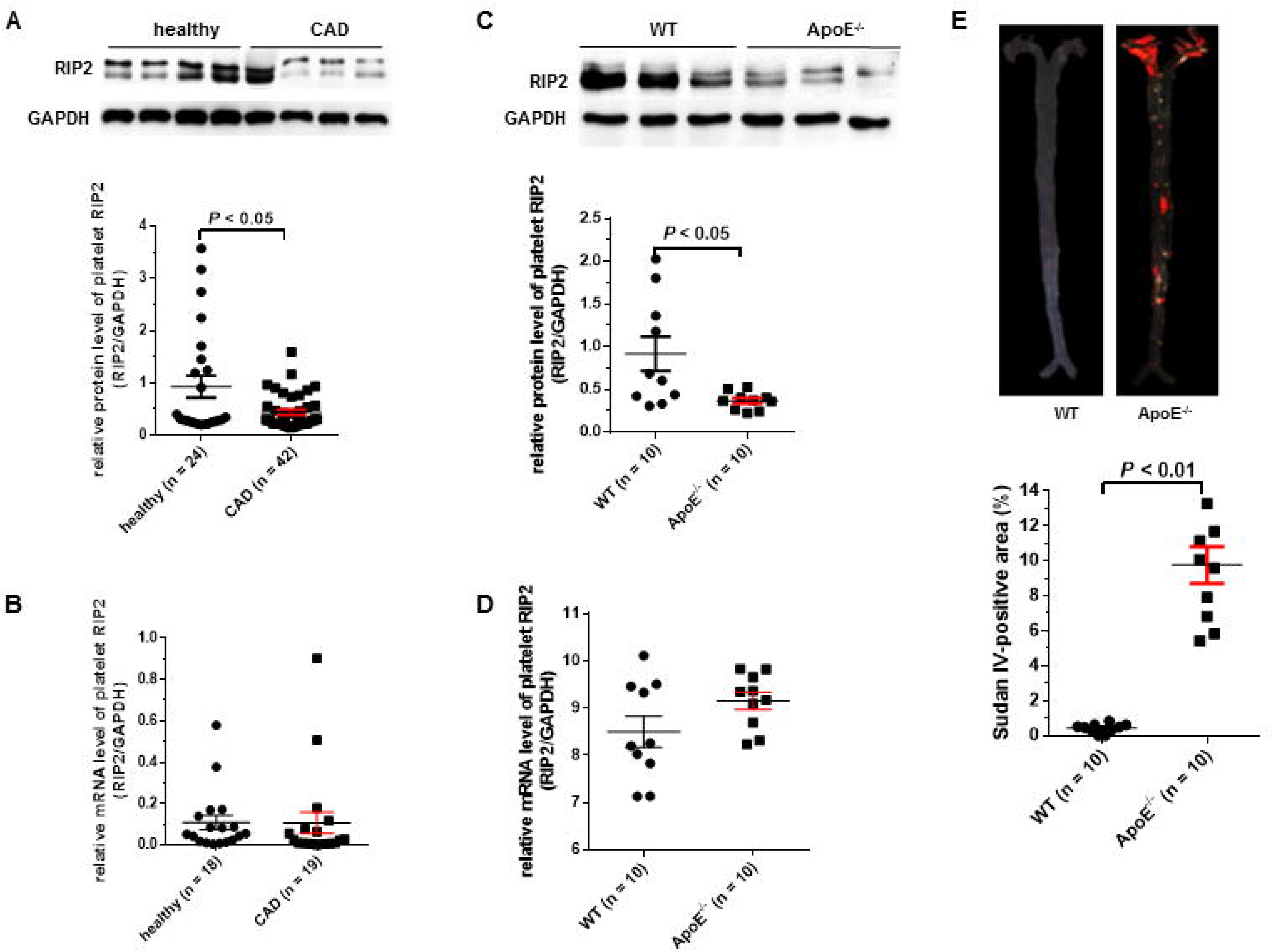
Impaired platelet RIP2 expression in patients with coronary artery heart disease and ApoE^-/-^ mice with atherosclerosis. **A.** Decreased RIP2 expression in platelets from patients with coronary artery heart disease (CAD) compared with healthy subjects analyzed by Western blot. Upper panel is the representative immunoblots of RIP2 and GAPDH from 4 healthy subjects and 4 patients; lower panel is the summary and statistical analysis of 24 healthy subjects and 42 patients. **B.** Similar RIP2 mRNA expression levels in platelets from CAD and healthy subjects analyzed by qPCR. **C.** Impaired platelet RIP2 expression in ApoE^-/-^ mice with atherosclerosis. Representative immunoblots from 3 ApoE^-/-^ with atherosclerosis and 3 wild type mice (upper panel) were presented, the summary (lower panel) was also provided. **D.** Similar RIP2 mRNA levels in platelets from ApoE^-/-^ mice with atherosclerosis and WT mice measured by qPCR. Data are expressed as mean ± SEM. **E**. Representative photographs showing exacerbated atherosclerosis injury in aorta from ApoE^-/-^ mice (upper panel) and the summary (lower panel) after high fat diet for 12 weeks compared with wild type mice receiving normal chow. Murine aorta was pinned out by *en face* technique and stained with Sudan IV.

### RIP2 deficient platelets deteriorate myocardial infarction

Atherosclerotic plaque rupture induces excessive platelet activation, occlusive thrombosis in coronary artery causing myocardial infarction. Timely reopening of the occluded vessels by percutaneous coronary intervention (PCI) or thrombolysis is the key to salvage the ischemic and still viable myocardium. However, the cardiac reperfusion itself following an acute myocardial infarction paradoxically generates ischemia/reperfusion (I/R) injury, further deteriorates myocardial damage, cardiac dysfunction, and death^39^. Among the multiple pathogenetic mechanisms underlying the I/R injury, platelets have emerged as a key player^39, 40^. By induction of myocardial infarction with I/R injury using platelet depletion/reconstitution model mice^27^, we found that platelet RIP2 deficiency significantly exacerbated I/R-induced cardiac contractile dysfunction in mice repopulated with RIP2^-/-^ platelets, as evaluated by ejection fraction (EF) and fractional shortening (FS) (Figure 8A). In line, the I/R-induced myocardial infarct size enhanced from 31.6% to 44.1% of the area at risk, although both groups had comparable areas at risk (Figure 8B). Thus, these *in vivo* data using the mouse model of myocardial infarction induced by I/R injury indicate that platelet RIP2 deficiency aggravates myocardial infarction and cardiac dysfunction.

**Figure 8.**
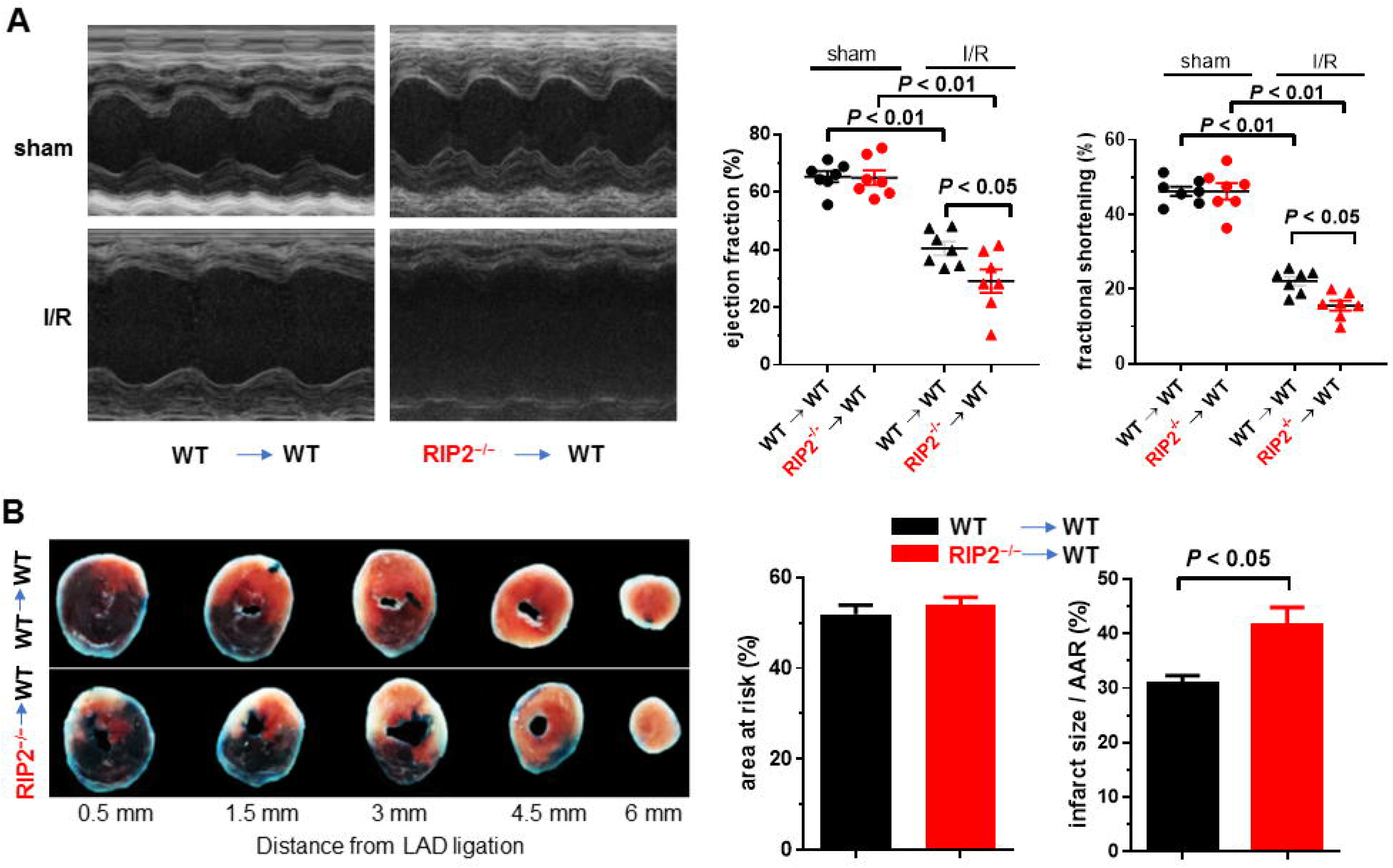
Platelet RIP2 deficiency aggravates myocardial function and infarction arear after ischemia/reperfusion (I/R) injury. **A.** Platelet RIP2 deficiency aggravates mouse myocardial function. Representative M-mode long-axis echo images and echocardiography quantification summary were provided. Each dot represents an individual animal. The results are expressed as mean ± SEM (n = 7). Two-way ANOVA followed by Tukey post hoc analysis was used for statistical analysis of myocardial. **B.** Platelet RIP2 deficiency aggravates mouse myocardial infarction. Representative infarction staining and the summary were provided. Student’s t-test was used to analyze infarct size. Platelet depletion/reconstitution model mice were subjected to I/R injury (45 min ischemia/24 hours reperfusion), M-mode echocardiography were taken and analyzed before hearts were isolated, sectioned, and stained. The echocardiography analysis and infarct size quantification were performed by investigators blind to the animal treatment.

## Discussion

In this study we found that RIP2 deficiency potentiated platelet activation, thrombosis, and myocardial infarction post I/R. The negative regulation of RIP2 on thrombosis and myocardial infarction is platelet specific, consistent with the reduced RIP2 expression in platelets from patients with coronary artery disease and mice with atherosclerosis. Mechanistically, we provided evidence that phosphorylated RIP2 reduces DOCK8-mediated Cdc42 activation and thus inhibits platelet dense granule secretion, independent of PRR activation. We also found that phosphorylated RIP2 associates with both GPIbα and filamin A. We therefore proposed that by strengthening GPIbα and filamin A binding, RIP2 keeps platelets in resting state and prevents overactivation when platelets are stimulated (Figure 6H).

Our results suggest that the negative regulation of RIP2 on platelet activation is GPIb-IX-V and GPVI pathway specific. This is reasonable, because GPVI physically and functionally associates with GPIb-IX-V, and GPIb-IX-V may act as an indirect collagen receptor using vWF as bridging molecule^41, 42^. GPVI is associated with GPIb-IX-V on platelet membrane, and a GPIbα-specific antibody inhibits platelet aggregation induced by collagen and CRP^43, 44^. Platelet glycoprotein V has also been shown to bind collagen and participate in platelet activation^45^. It appears that RIP2 negatively regulates platelet dense granule secretion primarily downstream of GPIb-IX-V.

Filamin A associates with GPIbα^46^. Englund et al reported that filamin A dissociation from GPIbα increases vWF binding to GPIbα^47^, suggesting a negative regulation of filamin A-GPIbα association on platelet activation. Consistently, we demonstrated significantly less GPIbα association with filamin A in platelets stimulated with thrombin and collagen in this study. Moreover, we show that RIP2 deficiency, which potentiates platelet activation, enhances the GPIbα dissociation from filamin A in platelets. We further reveal that phosphorylated RIP2 associates with both filamin A and GPIbα, inhibits platelet activation.

Cdc42 has been reported to regulate platelet dense granule secretion with contrarian results^35, 36^. Our findings that RIP2 deficiency potentiates dense granule release downstream of GPIbα and enhances Cdc42 activity support the positive regulation of Cdc42 on platelet dense granule secretion^36^. We hypothesize that Cdc42 mediates the negative regulation of RIP2 on platelet dense granule secretion. In support of this hypothesis, we found that DOCK8, the guanine nucleotide exchange factor for Cdc42 activation^38, 48^, is expressed in platelets and associates with p-RIP2. Moreover, RIP2 deficiency increases DOCK8 interaction with Cdc42 in platelets, in accordance with the enhanced Cdc42 activation in RIP2 deficient platelets. These results support that phosphorylated RIP2 sequesters DOCK8, prevents its binding to Cdc42 and the resultant activation of Cdc42, and inhibits platelet dense granule secretion downstream of GPIb-IX-V and GPVI pathway (Figure 6H).

Platelets play critical roles in both atherosclerosis and thrombotic complications^6^. Previously, Levin et al reported that RIP2 deficiency exacerbates atherosclerosis by increasing lipid accumulation in macrophages, suggesting the protective role of RIP2 against atherosclerosis^32^. This is in concert with our findings that RIP2 negatively regulates platelet activation and thrombosis. In line, we also found that platelets from patients with coronary artery disease and atherosclerotic mice express significantly less RIP2 protein. The impaired platelet RIP2 expression may thus dampen its protective effects against atherosclerosis, the resultant atherothrombosis, and myocardial infarction post I/R. The negative regulation of RIP2 on platelet dense granule secretion reported here may also contribute to the protective role of RIP2 against atherosclerosis reported by Levin et al^32^. Our findings are also in accordance with the recent report that platelet serotonin aggravates myocardial infarction^9^.

The mechanism underlying the reduced RIP2 protein in platelets from patients with coronary artery diseases and mice with atherosclerosis is not clear in the present study. Hyperlipidemia and inflammation are the two cornerstones synergizing the pathogenesis of atherosclerosis and atherothrombotic diseases^49^. Consistent with our findings, minimally modified LDL (mmLDL), a slightly oxidized LDL (oxLDL), which is significantly increased in the plasma of patients with coronary artery disease^50^, has been shown to reduce RIP2 in macrophages^32^. Infection and infection-induced inflammation are etiologically related to the pathogenesis of atherothrombotic diseases^51, 52^. Madrigal et al reported that lysine-specific bacterial cysteine protease of P. gingivalis (Kgp) induces the proteolysis of RIP2 in human aortic endothelial cells without influencing RIP2 mRNA^53^. Similar mechanisms may underlie the lower RIP2 protein levels in platelets from coronary artery disease patients and atherosclerotic mice in our study.

Using washed platelets assayed by light transmission aggregometry, though RIP2 deficiency enhances platelet dense release, we did not observe aggregation increase. Similar finding has also been reported by Fotinos et al^28^. When platelet aggregation was measured in whole blood using electric impedance aggregometry, we did observe the heightened platelet aggregation from RIP2^-/-^ mice. This discrepancy may be explained by the saturation of aggregatory response of light transmission aggregometry^54^, which does not always reflect platelet activation, especially when washed platelets were used^55^. On the other hand, white and red cells may also contribute to platelet aggregation regulated by RIP2, which may be better reflected by whole blood platelet aggregometry. Such findings also highlight the importance of simultaneous recording of aggregation and ATP release, because aggregation assay without platelet dense granule secretion measurement may miss the granule release abnormality in platelet function test using light transmission aggregometry.

Our study has some limitations. First, we reported RIP2 deficiency dense granule release without influence on α granule release; though similar findings have been reported by other group^29^, we did not go further to explore the mechanism, such as whether RIP2 affects other small GTPases associated with α granule release. Second, though the interaction of RIP2 with GPIb and filamin A explains the specific regulation of RIP2 on platelet activation downstream of GPIb and GPVI activation, how this interaction affects RIP2-DOCK8-Cdc42 pathway, and the resultant dense granule release is not clear. Interestingly, in line with our findings, GPIbα-filamin A interaction was suggested to be necessary for Cdc42 regulation downstream of GPIbα during platelets biogenesis^56^.

RIP2 was previously regarded as obligatory and specific mediator downstream of NOD action^57^. However, Johansson et al reported that myeloid-specific ablation of NOD2, but not its downstream kinase, RIP2, restrains the expansion of the lipid-rich necrotic core in Ldlr^-/-^ chimeric mice^58^, suggesting that NOD2 may function independent of RIP2. We previously reported that NOD2 activation potentiates platelet activation induced by thrombin and collogen, and NOD2 receptor agonist MDP induces RIP2 phosphorylation at serine 176, suggesting that RIP2 may mediate NOD2 signaling in platelets^18^. We found that RIP2 deficiency does not affect the potentiation effect of NOD2 activation on platelet aggregation induced by thrombin (data not shown), supporting that NOD2 may function in platelets RIP2 independently, consistent with the findings from Johansson et al^58^. Moreover, NOD2 deficiency does not impair platelet RIP2 phosphorylation (serine 176) induced by thrombin and collagen (Online-only Data supplement Figure IV), further validating the PRR-independent RIP2 activation in platelets.

In conclusion, through a novel PRR-independent pathway, p-RIP2-DOCK8-Cdc42, RIP2 inhibits platelet activation, thrombosis, and ameliorates myocardial infarction post I/R. Activating platelet RIP2 pathway may be a promising approach for therapeutic intervention of atherothrombotic diseases from the early atherosclerosis stage to coronary events.

## Acknowledgments

JZ designed and performed the research, collected and analyzed data, and wrote the paper. GP, YL, LC, LH, ZQ, ZZ, YZ and SZ performed experiments and helped to analyze data. JQ provided materials and contributed to the writing of the manuscript. XX and KD analyzed data and contributed to the writing. ZD initiated and supervised the project, designed research, analyzed, and interpreted results, and wrote the manuscript. We are grateful to Dr. Taei Matsui from Fujta Health University School of Health Sciences to provide botrocetin. The authors thank Dr. Satya P. Kunapuli from Temple University Sol Sherry Thrombosis Research Center and Dr. Hu Hu from Zhejiang University School of Medicine for critically reading the manuscript.

## Disclosures

None.

**Supplemental Materials**

Supplementary Tables I - IV

Supplementary Figures I – IV

Expanded Materials and Methods

## Abbreviations

RIP2: Receptor-interacting protein 2
PRRs: pattern recognition receptors
MI: myocardial infarction
GPIb: Platelet glycoprotein Ib
GPVI: Platelet glycoprotein VI
DOCK8: dedicator of cytogenesis protein 8
Cdc42: cell division cycle 42
TXA2: thromboxane A2
vWF: von Willebrand factor
NOD2: Nucleotide-binding oligomerization domain 2
CARD: Caspase Recruitment Domain
CRP: collagen related peptide
CAD: coronary heart disease

